# Protection of catalytic cofactors by polypeptides as a driver for the emergence of primordial enzymes

**DOI:** 10.1101/2023.03.14.532612

**Authors:** Luis I. Gutierrez-Rus, Gloria Gamiz-Arco, J.A. Gavira, Eric A. Gaucher, Valeria A. Risso, Jose M. Sanchez-Ruiz

## Abstract

Enzymes catalyze the chemical reactions of life. For nearly half of known enzymes, catalysis requires the binding of small molecules known as cofactors. Polypeptide-cofactor complexes likely formed at a primordial stage and became starting points for the evolution of many efficient enzymes. Yet, evolution has no foresight so the driver for the primordial complex formation is unknown. Here, we use a resurrected ancestral TIM-barrel protein to identify one potential driver. Heme binding at a flexible region of the ancestral structure yields a peroxidation catalyst with enhanced efficiency when compared to free heme. This enhancement, however, does not arise from protein-mediated promotion of catalysis. Rather, it reflects protection of bound heme from common degradation processes and a resulting longer life time and higher effective concentration for the catalyst. Protection of catalytic cofactors by polypeptides emerges as a general mechanism to enhance catalysis and may have plausibly benefited primordial polypeptide-cofactor associations.

## Introduction

Life involves a vast network of chemical reactions, most of which would not occur at a sufficient rate in the absence of effective enzyme catalysis. With the obvious exception of ribozymes, enzymes are based on protein scaffolds. Most, if not all, of modern enzyme activities have evolved from prior enzyme activities (Ohno 1970; Khersonsky and Tawfik 2010). This evolutionary process is reasonably well-understood and protein engineers can mimic it in the lab using directed evolution (Campbell et al 2016; Zeymer and Hilvert 2018). On the other hand, it is a logical necessity that the first enzymes did not arise from previously existing “older” enzymes. That is, the first primordial enzymes must have necessarily arisen *de novo* in previously non-catalytic protein scaffolds. Reconstructions of the gene content of the last universal common ancestor (LUCA) support that a diversity of enzymes already catalyzed many different reactions at a very early evolutionary stage (Weiss et al 2016). It seems reasonable, therefore, that there are efficient mechanisms for the *de novo* emergence of completely new enzymes. Such mechanisms, however, remain elusive to protein engineers, who have found the generation of efficient *de novo* enzymes in the lab to be extremely challenging (Blomberg et al 2013; Risso et al 2017; Donnelly et al 2018; Lovelock et al 2022; Yeh et al 2023). Our limited grasp of the evolutionary mechanisms of *de novo* enzyme generation implies a serious gap in our understanding of the origin of life.

A substantial fraction (around 50%) of enzymes rely on cofactors for catalysis (Fischer et al 2010). This statement is true for many modern enzymes, but it also very likely holds for the most ancient enzymes, as indicated by reconstructions of the gene content of LUCA (Weiss et al 2016). Many cofactors, in particular metals and metal-containing organic cofactors, display by themselves significant levels of catalysis with a diversity of chemical reactions (Andreini et al 2008; Fischer et al 2010). Both polypeptides and cofactors were likely available already at a prebiotic stage (Frenkel-Pinter et al 2020; Goldman and Kacar 2021). Therefore, recruitment of catalytic cofactors by polypeptides would appear to provide a simple and straightforward mechanism for the *de novo* emergence of many of the primordial enzymes (although certainly not of all of them). On closer examination, however, this mechanism has a serious shortcoming: the driving force for the recruitment is not apparent at all. Certainly, once a polypeptide-cofactor complex has been formed, new possibilities arise in a Darwinian evolution scenario, including the enhancement of catalysis through mutations in the protein moiety. Yet, we know evolution has no foresight (Jacob 1977), so any driver of primordial polypeptide-cofactor association must have provided an immediate selective advantage, such as an instant enhancement in catalysis. However, the opposite would seem reasonable on general grounds. Association of a cofactor with a “naïve” polypeptide, *i*.*e*., a polypeptide that has not evolved to enhance catalysis by the cofactor, would simply decrease cofactor exposure to the environment and therefore its accessibility to substrates, thus likely impairing catalysis.

Here, we report experimental studies on the emergence of cofactor-based catalysis in an ancestral TIM-barrel protein. Our experiments reveal a general mechanism of immediate catalysis enhancement that may have plausibly provided a driving force for the polypeptide-cofactor association during a primordial period.

## Results

The TIM-barrel is the most widespread enzyme fold, providing a structural scaffold for a large diversity of modern enzyme functionalities (Wierenga 2001; Nagano et al 2002). In fact, the TIM-barrel fold has been proposed to have played an essential role in early metabolism (Goldman et al 2016). We recently reported an ancestral reconstruction exercise targeting family 1 glycosidases, which are of the TIM-barrel fold (Gamiz-Arco et al 2021). We found a putative common ancestor of eukaryotic and bacterial family 1 glycosidases that displayed unusual properties, including tight and stoichiometric binding of heme at a conformationally flexible region distinct from the glycosidase active site (figs. 1A, 1B and 1C). Remarkably, heme binding is extremely rare among modern TIM-barrel proteins (Gamiz-Arco et al 2021).

**Fig. 1.**
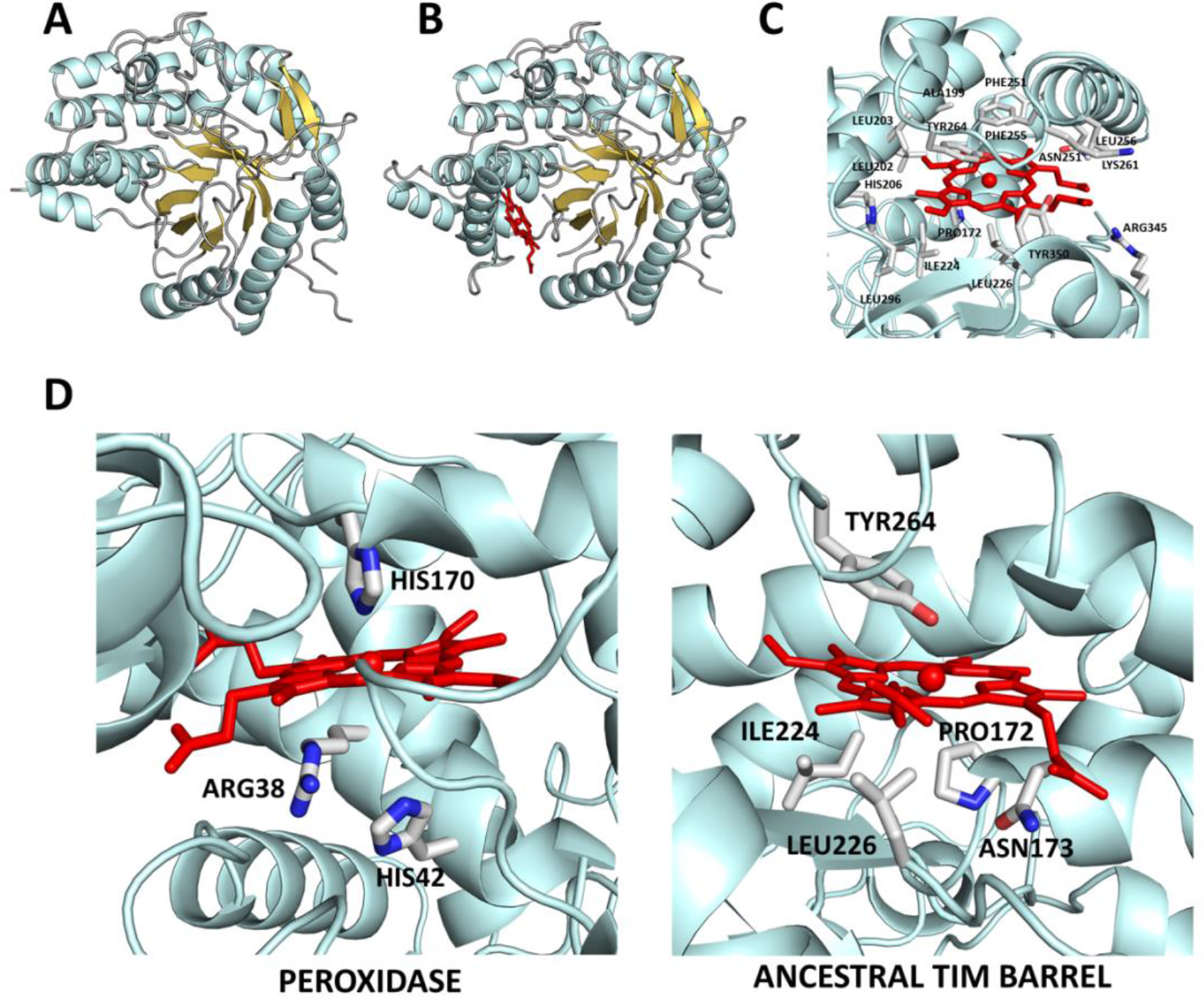
3D-structure of the ancestral TIM-barrel glycosidase studied in this work as determined by X-ray crystallography (Gamiz-Arco et al 2021). (A) Structure of the protein without heme bound (PDB ID 6Z1H). Note that the region where heme binds displays substantial conformational flexibility and that, as a result, there are missing sections in the electronic density maps (for further details, see Gamiz-Arco et al 2021). (B) Structure of the protein with heme bound (PDB ID 6Z1M). Heme binding rigidifies the protein and allows a larger part of the structure to be determined from the density maps, as it is apparent by comparing the structure shown in B with that shown in A (for further details, see Gamiz-Arco et al. 2021). (C) Zoom-in of the heme-binding region providing amino acid detail. (D) Comparison of the molecular environments of the heme in horseradish peroxidase (PDB ID 1HCH) and the ancestral TIM-barrel (PDB ID 6Z1M). Critical residues for peroxidase catalysis (the proximal histidine and the distal histidine and arginine) are highlighted in the peroxidase structure. In the ancestral TIM-barrel structure, the proximal residue is not a histidine and the residues at the opposite side of the heme ring (labeled) do not include histidines or arginines.

Free heme is known to display a low but measurable peroxidase activity (Brown et al 1969). Therefore, heme binding may be expected to confer some peroxidase activity to the ancestral TIM-barrel. We have used the common peroxidase substrate o-dianisidine (fig. 2A) to compare the peroxidase activity of heme with that of heme bound to the ancestral scaffold. Peroxidation of o-dianisidine leads to absorption in the visible region of the spectrum, which that can be used to follow the chemical reaction (Jenkins et al 2021). As it is customary in enzyme kinetics studies, we started with determinations of the initial rates of the reaction, which can be easily calculated from the initial increases of the absorbance due to the peroxidation product (fig. 2B). We performed a large number of experiments in solutions at different pH values. In most of the studied pH range, the peroxidase activity of the free heme is substantially higher than that of the heme-bound ancestral scaffold (fig. 2B). This result is not surprising, given that burial of the heme in the ancestral protein structure (see figure 7C in Gamiz-Arco et al 2021) should hinder the substrate access to the cofactor, thus impairing catalysis. Furthermore, the heme binding region of the ancestral structure has not evolved to promote cofactor-based catalysis. Enhancement of cofactor-based catalysis in modern peroxidase enzymes is linked to a specific molecular machinery that includes the proximal histidine and distal histidine and arginine residues (Poulos and Kraut 1980; Ortiz de Montellano 2010). Such assistance molecular machinery is lacking from the ancestral TIM-barrel (fig. 1D).

**Fig. 2.**
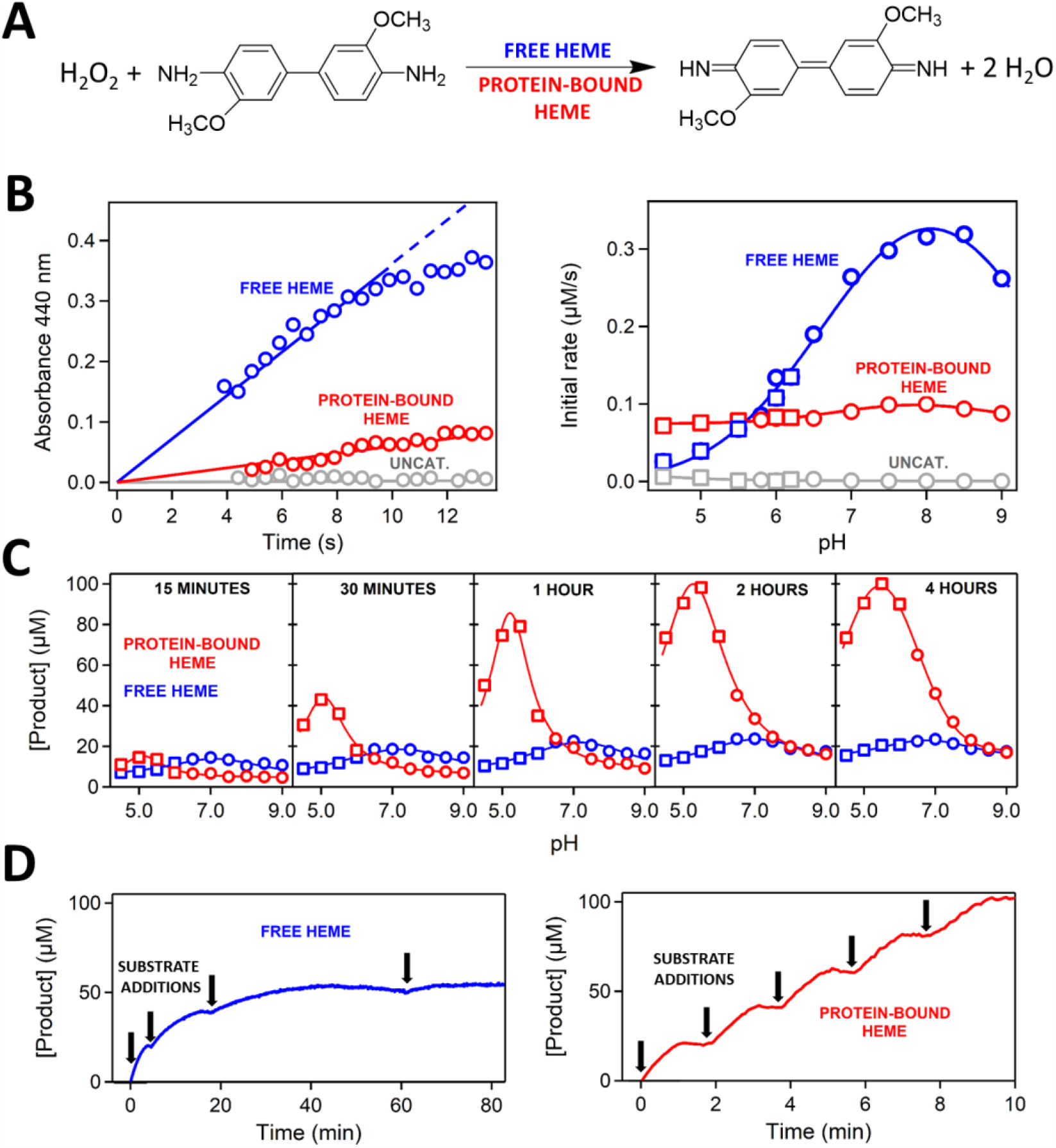
Peroxidase activity of free heme versus heme bound to the ancestral TIM-barrel. (A) Reaction used to test peroxidase activity. The kinetics of o-dianisidine peroxidation can be determined by following the increase of absorbance at 440 nm upon peroxidation. (B) Left: representative examples of the profiles of absorbance at 440 nm versus time used to calculate the initial rates of the reaction. The example shown corresponds to pH 7, heme concentration of 0.4 μM (either free or protein bound) and initial concentrations of hydrogen peroxide and o-dianisidine of 10 mM and 0.1 mM, respectively. The data show that the initial reaction rate is higher with free heme as catalyst. Right: plot of initial rate versus pH for free heme, protein-bound heme and a control experiment in the absence of heme (labeled “uncat”). In most of the studied pH range, initial rates are higher with free heme as catalyst. Heme and initial substrate concentrations are the same as in B. Symbols refer to the buffer used: squares, 200 mM acetate, 150 mM NaCl; circles, 200 mM phosphate, 150 mM NaCl. (C) Profiles of amount of peroxidation product formed versus pH over long reaction times (shown inside the plots). As the reaction time increases, bound heme progressively becomes a more efficient peroxidation catalyst than free heme. Buffers, heme concentration and initial substrate concentrations are the same as in B. (D) Peroxidation kinetics over long reaction times with free heme (left) and protein-bound heme (right) as catalysts with repeated additions of 0.1 mM o-dianisidine substrate (labeled with arrows). Degradation of free heme catalysis is apparent by the progressive absence of peroxidation of freshly added o-dianisine. By contrast, no such degradation is observed with protein-bound heme. The data correspond to pH 7, heme concentration of 2 μM (either free or protein bound) and initial concentration of hydrogen peroxide of 10 mM.

Subsequently, we determined the total yield of peroxidation product for several pH values at times ranging from 15 minutes to 4 hours. Strikingly (fig. 2C) we found a much higher conversion of substrate to product with the heme-bound ancestor when compared to free heme, in particular for reaction times on the order of hours. A reasonable explanation for these results is that free heme undergoes time-dependent processes that impair its peroxidase activity, while heme bound to the protein is protected and retains its peroxidase activity during a much longer time. To test this hypothesis, we performed experiments at longer durations and with repeated additions of substrate (fig. 2D). We found that, unlike the heme bound to the ancestral protein, free heme gradually loses its capability to peroxidize the freshly added substrate.

To further explore the mechanisms of heme protection upon binding to the ancestral protein, we performed kinetic experiments in which heme, either free or protein-bound, was incubated for given times in the reaction buffer prior to substrate addition. Profiles of product concentration versus reaction time for these experiments are shown in fig. 3A. Interestingly, we found the final substrate yield of reaction catalyzed by free heme to decrease with incubation time. This decrease is most likely linked to heme self-association during incubation time and the consequent reduction in the concentration of the more active heme monomer. In support of this interpretation, the alteration in the Soret band known to be linked to heme self-association (Inada and Shibata 1962) occurs in the same time scale as does the decrease in peroxidase activity observed upon incubation (figs. 3B and 3C). Heme bound to the ancestral protein cannot associate with other heme molecules and it is therefore protected from the loss of peroxidase activity caused by self-association.

**Fig 3.**
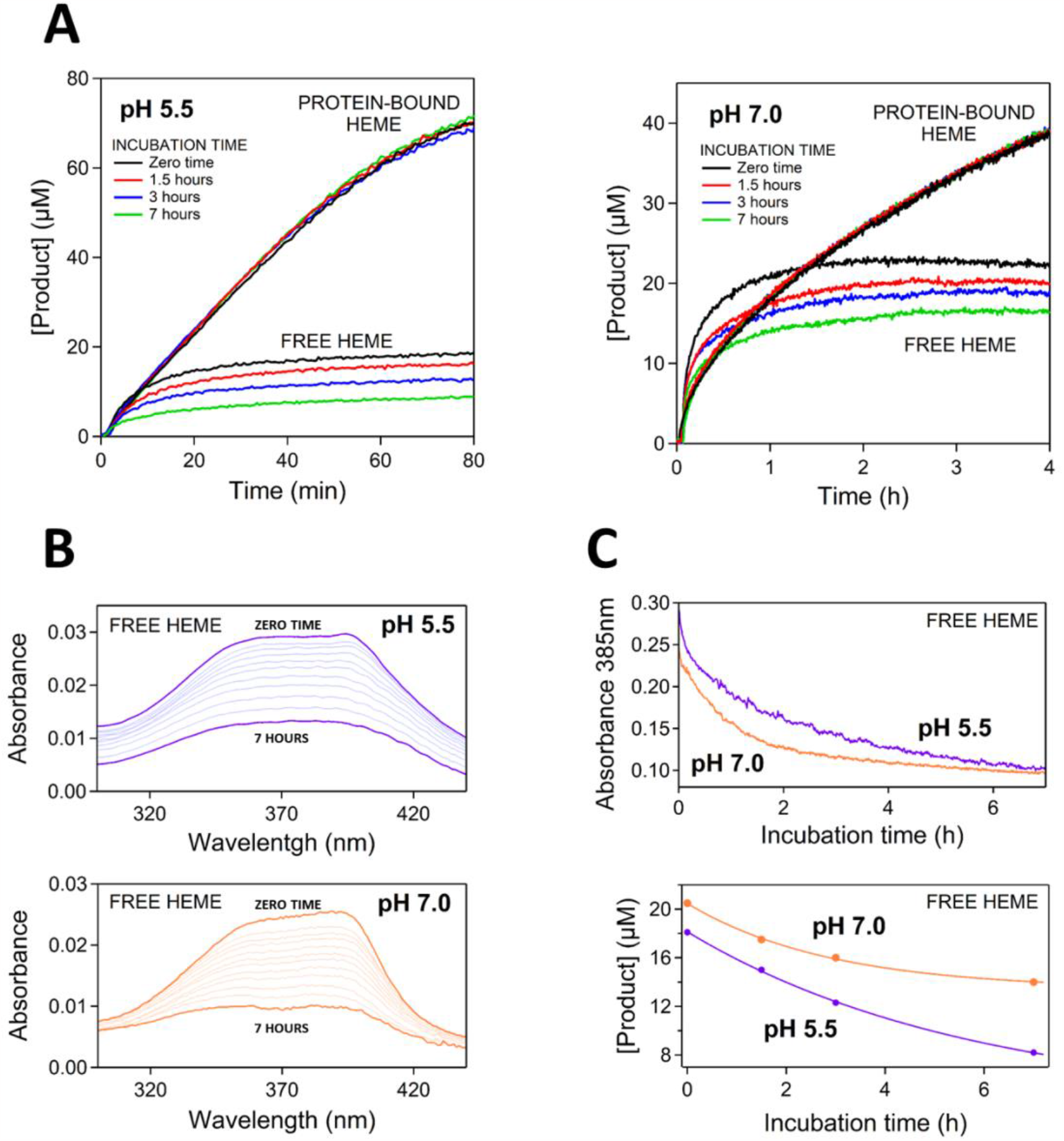
Degradation of catalysis by free heme during and before the peroxidation reaction. (A) Peroxidation kinetics over long reaction times with free heme and protein-bound heme as catalysts for pH 5.5 and pH 7. In all experiments, the heme concentration (either free protein-bound) was 0.4 μM and the initial concentrations of o-dianisidine and hydrogen peroxide were 0.1 mM and 10 mM, respectively. The experiments differ in the incubation duration, *i*.*e*., the time free heme or protein-bound heme were kept in the solution before addition of the substrates. The curvature of the kinetic profiles for free heme reveal severe catalysis degradation during the peroxidation reaction. Yet, substantial degradation also occurs during the incubation time as shown by decreasing final product yields with increasing incubation time. (B) Self-association of heme as revealed by flattening of the Soret band of free heme in solution. Spectra were collected approximately every 30 minutes from the dilution of the heme stock in the buffer (zero time) to seven hours. See legend to fig. 2 and Methods for buffer composition. (C) Plots of absorbance at 385 nm versus time (from the data given in B) and plots of final product yield versus incubation time (from the data given in A for free heme) Self-association occurs in the same time scale as (and it is the likely cause of) the degradation of catalytic potential of free heme during the incubation time.

It is also clear from the strong curvature and the leveling-off of the kinetic profiles at long reaction times (fig 3A and fig. 2D) that the degradation of the peroxidase activity of free heme is substantially accelerated after substrate addition. This points to an additional mechanism of inactivation acting during the catalytic cycle, likely involving chemical alterations of the heme caused by reactive oxygen species, as it has been described for heme in peroxidases (Valderrama et al 2002) and for free heme in solution (Brown et al 1968; Brown and Jones 1968). Remarkably, the loss of peroxidase activity during the catalytic cycle appears to be considerably limited in the restricted and defined molecular environment provided by the ancestral TIM-barrel for the bound heme (figs. 3A and 2D).

## Discussion

Heme binding at a conformationally flexible region of an ancestral TIM-barrel leads to an enhancement in the efficiency of heme as a peroxidation catalyst, as shown by the increase in peroxidation reaction yield over a time scale of hours. Determination of initial reaction rates, however, indicates that binding does not improve the intrinsic peroxidase activity of heme. In fact, the opposite is true within most of the studied pH range: initial reaction rates are somewhat lower with the protein-heme complex as compared with free heme, likely reflecting limited substrate access to the substantially-buried bound heme. Therefore, the enhancement in the efficiency of peroxidation catalysis upon heme binding to the protein scaffold cannot be attributed to promotion of cofactor catalysis by the protein moiety, as is the case with modern peroxidases (fig. 1D). In fact, our experimental data (figs. 2 and 3) clearly support that the enhancement is due to the retardation of processes that impair the peroxidase activity of heme, resulting in a much longer life time and a higher effective concentration for the catalyst when it is bound to the protein.

The ancestral TIM-barrel we have used as protein scaffold in our experiments is a putative ancestor of bacterial and eukaryotic family-1 glycosidases (Gamiz-Arco et al 2021) and may perhaps probe an early stage in glycosidase evolution. However, in the context of the evolutionary history of proteins, it belongs to a period long after the generation of the first enzymes. Yet, the consequences for catalysis of heme binding to a conformationally flexible region of the ancestral TIM-barrel point to and mimic a plausible scenario for the primordial emergence of enzymes. We briefly elaborate this scenario below.

Reasonable chemical mechanisms for the prebiotic formation of polypeptides have been proposed (Frenkel-Pinter et al 2020). Indeed, it has been hypothesized that polypeptides already existed in the primordial RNA world, where they served as enhancers of ribozyme activity (Romero et al, 2016; Wolf and Koonin 2017). Regarding protein cofactors, their extreme evolutionary conservation strongly suggests that they are very ancient (Chu and Zhang 2020). The likely origin of inorganic cofactors in the geochemical environment in which life begun and that of organic cofactors (coenzymes) as parts of primordial ribozymes have been noted (Goldman and Kacar 2021). Overall, there can be little doubt that polypeptides and cofactors co-existed at some primordial stage. Then, their association may have immediately promoted catalysis by increasing the life time and the effective concentration of catalytic cofactors. It follows that, as soon as some means of transmission of genetic information, even if rudimentary and error-prone, was available, natural selection for polypeptides with increasing capability to protect catalytic cofactors became possible. This selection could have also favored cofactors that interact efficiently with polypeptides and become readily protected. Lastly, the stable and catalytically competent polypeptide-cofactors complexes thus formed provided starting points for the Darwinian evolution of a protein molecular machinery that assisted and further enhanced cofactor-based catalysis.

The protection mechanism for catalysis enhancement has been demonstrated in this work on the basis of the peroxidase activity of heme, but it should apply to many other cofactors. The emergence of enzymes based on catalytic iron-sulfur clusters may be a particularly relevant example. Iron and sulfur were abundant in the geochemical environment that likely hosted primordial life (Beinert 2000) and enzymes based on iron-sulfur clusters abound in reconstructions of the gene content of LUCA (Weiss et al 2016). Free iron-sulfur clusters have been found to mimic the basic features of protein-bound clusters, but in non-aqueous media and in the absence oxygen (Beinert 2000). Even if destruction by reaction with oxygen was not an issue in an anaerobic primordial environment (Weiss et al 2016), iron-sulfur clusters appear as obvious candidates for primordial catalysis enhancement through protection by polypeptides (Kim et al 2018).

Ferredoxins are iron-sulfur proteins that perform electron transfer in a diversity of biochemical transformations. About 60 years ago, Margaret Dayhoff noted the simplicity of ferredoxins, which consist of an inorganic active site and a very short polypeptide which she proposed to have emerged by duplication of smaller peptides (Eck and Dayhoff 1966). Modern ferredoxins may, therefore, be relics of primordial protection events. An intriguing, albeit speculative, possibility is that heme binding to our ancestral TIM-barrel glycosidase is also a relic of primordial protection, although, in this case, the ancestral functional feature has undergone evolutionary degradation and it is only observed in vestigial form in modern glycosidases (Gamiz-Arco et al 2021).

## Methods

The ancestral TIM-barrel glycosidase was prepared as previously described (Gamiz-Arco et al., 2021). Briefly, the gene for the His-tagged protein in a pET24 vector was cloned into *E. coli* BL21 (DE3) cells and the protein was purified using Ni-NTA chromatography. Protein prepared in this way typically has a very small amount of bound heme. Ancestral protein saturated with heme was prepared by incubation with 5-fold excess of heme followed by size-exclusion chromatography to eliminate non-bound heme, as we have previously described in detail (Gamiz-Arco et al 2021). The heme to protein ratio was found to be close to unity on the basis of absorbance determinations for the protein band at 280 nm and the heme Soret band.

As previously described (Gamiz-Arco et al., 2021), heme solutions were prepared by high-dilution (typically 1:1000) in the desired buffer of a stock solution in concentrated sodium hydroxide and used immediately. Stock solutions of heme were prepared daily. Heme concentrations in stock solution were determined from the absorbance of the heme Soret band at 385 nm using a known value of the extinction coefficient (Deniau et al 2003). Stock solutions of o-dianisidine were prepared by weight. Stock solutions of hydrogen peroxide were prepared by dilution of commercially available stock solution and their concentrations were determined from the absorbance at 240 nm using a known extinction coefficient (Jiang et al 1990). The peroxidation reaction was initiated by adding microliter volumes of the reactants to a 2-mL solution containing free heme or protein-bound heme. The reaction was followed by measuring the absorbance of the peroxidation product at 440 nm. A known value of the extinction coefficient (Jenkins et al 2021) was used calculate substrate concentration from absorbance values. Peroxidation experiments were performed in wide pH range using the following buffers: 200 mM acetate, 150 mM NaCl for the pH range 4.5-6.2 and 200 mM phosphate, 150 mM NaCl for the pH range 5.8-9.0.

## Acknowledgments

This work was supported by Human Frontier Science Program grant RGP0041/2017 (J.M.S.R. and E.A.G.), National Science Foundation grant 2032315 (E.A.G.), Department of Defense grant MURI W911NF-16-1-0372 (E.A.G.), National Institutes of Health grant R01AR069137 (E.A.G.), Spanish Ministry of Science and Innovation / FEDER Funds grant PID2021-124534OB-100 (J.M.S.R.) and grant PID2020-116261GB-I00 (J.A.G.).

## Author Contributions

L.I.G.R. carried out protein preparation, and designed and performed the experimental determination peroxidase activity under the supervision of V.A.R.; G.G.A. provided essential input regarding the catalytic properties of the ancestral glycosidase; J.A.G. provided essential input regarding the structural interpretation of the protection mechanisms; E.A.G provided essential input regarding the interpretation of the data in an evolutionary context; J.M.S.R. designed the research and wrote the first draft of the manuscript; all authors discussed the manuscript, suggested modifications and improvements and contributed to the final version.

## Data Availability

All relevant experimental data are provided in the main-text figures. Tables with the data are available upon reasonable request.

## Conflict of interest statement

The authors declare no competing interests.

## Notes

### Competing Interest Statement

The authors have declared no competing interest.

## References

Andreini C, Bertini I, Cavallaro G, Holliday GL, Thornton JM. 2008. Metal ions in biological catalysis: from enzyme databases to general principles. J Biol Inorg Chem 13:1205–1218.

Beinert H. 2000. Iron-sulfur proteins: ancient structures, still full of surprises. J Biol Inorg Chem 5:2–15.

Blomberg R, Kries H, Pinkas DM, Mittl PRE, Grütter MG, Privett HK, Mayo SL, Hilvert D. 2013. Precision is essential for efficient catalysis in an evolved Kemp eliminase. Nature 503:418–421.

Brown SB, Jones P, Suggett. 1968. Reactions between haemin and hydrogen peroxide. Part1.-Ageing and non-destructuve oxidation of haemin. Transactions of the Faraday Society 64:986–993.

Brown SB, Jones P. 1968. Reactions between haemin and hydrogen peroxide. Part 2.-Destructive oxidation of heamin. Transactions of the Faraday Society 64:994–998.

Brown SB, Dean TC, Jones P, Kremer ML. Catalytic activity of haemin. 1970. Transactions of the Faraday Society 66:1485–1490.

Campbell E, Kaltenbach M, Correy GJ, Carr OD, Porebski BT, Livingstone EK, Afriat-Jurnow L, Buckle AM, Weik M, Hollfelder F, Tokuriki N, Jackson CJ. 2016. The role of protein dynamics in the evolution of new enzyme function. Nat Chem Biol 12:944–950.

Chu XY, Zhang HY. 2020. Cofactors as molecular fossils to trace the origin and evolution of proteins. ChemBioChem 21:3161–3168.

Deniau C, Gilli R, Izadi-Pruneyre N, Létoffé Delepierre M, Wandersman C, Briand C, Lecroisey A. 2003. Thermodynamics of heme binding to the HasA_SM_hemophore: effect of mutations and three key residues for heme update. Biochemistry 42:10627–10633.

Donnelly AE, Murphy GS, Digianantonio KM, Hecht MH. 2018. A de novo enzyme caalyzes a life-sustaining reaction in Escherichia coli. Nat Chem Biol 14:253–255.

Eck RV, Dayhoff MO. 1966. Evolution of the structure of ferredoxin based on living relics of primitive amino acid sequences. Science 152:363–366.

Fischer JD, Holliday GL, Rahman SA, Thornton JM. 2010. The structures and physicochemical properties of organic cofactors in biocatalysis. J Mol Biol 403:803–824.

Frenkel-Pinter M, Samanta M, Ashkenasy G, Leman LJ. 2020. Prebiotic peptides: molecular hubs in the origin of life. Chem Rev 120:4707–4765.

Gamiz-Arco G, Gutierrez-Ruz LI, Risso VA, Ibarra-Molero B, Hoshino Y, Petrovic D, Justicia J, Cuerva JM, Romero-Rivera A, Seelig B, Gavira JA, Kamerlin SCL, Gaucher EA, Sanchez-Ruiz JM. 2021. Heme-binding enables allosteric modulation in an ancient TIM-barrel glycosidase. Nat Comunn 2:380.

Goldman AD, Beatty JT, Landweber LF. 2016. The TIM barrel architecture facilitated the early evolution of protein-mediated metabolism. J Mol Evol 82:17–26.

Goldman AD, Kacar B. 2021. Cofactors are remnants of life’s origin and early evolution. J Mol Evol 89:127–133.

Inada Y, Shibata K. 1962. Soret band of monomeric hematin and its changes on polymerization. Biochem Biophys Res Commun 9:323–327.

Jacob F. 1977. Evolution and tinkering. Science 196:1161–1166.

Jenkins JMX, Noble CEM, Grayson KJ, Mulholland AJ, Anderson R. 2021. Substrate promiscuity of a de novo designed peroxidase. J Inorg Chem 217:111370.

Jiang ZY, Woollard ACS, Wolff SP. 1990. Hydrogen peroxide production during experimental protein glycation. FEBS Lett 268:69–71.

Khersonsky O, Tawfik DS. 2010. Enzyme promiscuity: a mechanistic and evolutioanary perspective. Annu Rev Biochem 79:471–505.

Kim JD, Pike DH, Tyryshkin AM, Swampa GVT, Raanan H, Montelione GT, Nanda V, Falkowski PG. 2018. Minimal heterochiral de novo designed 4Fe-4S binding peptide capable of robust electron transfer. J Am Chem Soc 140:11210–11213.

Lovelock SL, Crawshaw R, Basler S, Levy C, Baker D, Hilvert D, Green AP. 2022. The road to fully programmable protein catalysis. Nature 606:49–58.

Nagano N, Orengo CA, Thornton JM. 2002. One fold with many functions: the evolutionary relationships between TIM barrel families based on their sequences, structures and functions. J Mol Biol 321:741–765.

Ohno S. 1970. Evolution by gene duplication. Berlin, Germany, Springer.

Ortiz de Montellano PR. 2010. Catalytic mechanisms of heme peroxidases. In E. Torres, M Ayala (eds.) “Biochatalysis based on heme peroxidases” p 79–107. New York, USA, Springer.

Poulos TL, Kraut J. 1980. The stereochemistry of peroxidase catalysts. J Biol Chem 255:8199–8205.

Risso VA, Martinez-Rodriguez S, Candel AM, Krüger DM, Pantoja-Uceda D, Ortega-Muñoz M, Santoyo-Gonzalez F, Gaucher EA, Kamerlin SCL, Bruix M, Gavira JA, Sanchez-Ruiz JM. De novo active sites for resurrected Precambrian enzymes. Nat Commun 8:16113.

Romero MLR, Rabin A, Tawfik DS. 2016. Functional proteins from short peptides: Dayhoff’s hypothesis turns 50. Angew Chem Int Ed 55:15966–15971.

Valderrama B, Ayala M, Vazquez-Duhalt R. 2002. Suicide inactivation of peroxidases and the challenge of engineering more robust enzymes. Chem Biol 9:555–565.

Weiss MC, Soussa FL, Mrnjavac N, Neukirchen S, Roettger M, Nelson-Sathi S, Martin WF. 2016. The physiology and habitat of the last universal common ancestor. Nat Microbiol 1:16116.

Wierenga RK. 2001. The TIM-barrel fold: a versatile framework for efficient enzymes. FEBS Lett 492:193–198.

Wolf YI, Koonin EV. 2007. On the origin of the translation system and the genetic code in the RNA world by means of natural selection, exaptation, and subfunctionalization. Biol Direct 2:14.

Yeh AHW, Norn C, Kipnis Y, Tischer D, Pellock SJ, Evans, D, Ma P, Lee GR, Zhang JZ, Anishchenko I, Coventry B, Cao L, Dauparas J, Halabiya S, DeWitt M, Carter L, Houk KN, Baker D. 2023. De novo design of luciferases using deep learning. Nature 614:774–780.

Zeymer C, Hilvert D. 2018. Directed evolution of protein catalysts. Annu Rev Biochem 87:131–157.

